# ODL-BCI: Optimal deep learning model for brain-computer interface to classify students confusion via hyperparameter tuning

**DOI:** 10.1101/2023.10.30.564829

**Authors:** Md Ochiuddin Miah, Umme Habiba, Md Faisal Kabir

## Abstract

Brain-computer interface (BCI) research has gained increasing attention in educational contexts, offering the potential to monitor and enhance students’ cognitive states. Real-time classification of students’ confusion levels using electroencephalogram (EEG) data presents a significant challenge in this domain. Since real-time EEG data is dynamic and highly dimensional, current approaches have some limitations for predicting mental states based on this data. This paper introduces an optimal deep learning (DL) model for the BCI, ODL-BCI, optimized through hyperparameter tuning techniques to address the limitations of classifying students’ confusion in real time. Leveraging the “confused student EEG brainwave” dataset, we employ Bayesian optimization to fine-tune hyperparameters of the proposed DL model. The model architecture comprises input and output layers, with several hidden layers whose nodes, activation functions, and learning rates are determined utilizing selected hyperparameters. We evaluate and compare the proposed model with some state-of-the-art methods and standard machine learning (ML) classifiers, including Decision Tree, AdaBoost, Bagging, MLP, Näıve Bayes, Random Forest, SVM, and XG Boost, on the EEG confusion dataset. Our experimental results demonstrate the superiority of the optimized DL model, ODL-BCI. It boosts the accuracy between 4% and 9% over the current approaches, outperforming all other classifiers in the process. The ODL-BCI implementation source codes can be accessed by anyone at https://github.com/MdOchiuddinMiah/ODL-BCI.

## 1. Introduction

The evolution of BCI technology has unlocked groundbreaking pathways in cognitive neuroscience, especially in understanding and interpreting EEG data [1, 2, 3]. EEG, a representation of the brain’s electrical activity, offers invaluable insights into cognitive states and has diverse applications, ranging from medical diagnostics to enhancing learning experiences in education [4, 5, 6]. One critical aspect lies in monitoring and assessing students’ confusion levels in education, which can provide valuable insights into their learning behaviours and enable educators to tailor personalized teaching methods for improved learning outcomes [7, 8]. Despite its potential, extracting meaningful information from EEG data is complex, requiring sophisticated analytical models to handle its high dimensionality and variability [9, 10].

Recent advancements in deep learning have shown exceptional promise in analyzing EEG signals, providing the ability to learn intricate patterns and associations from vast and complex datasets [11]. Nevertheless, the performance of DL models is heavily contingent upon the careful selection of hyperparameters [12]. Manual tuning is not only laborious and time-consuming but also rarely yields optimal results [13]. This has led researchers to seek automated, intelligent methods for hyperparameter optimization [14].

However, the right choice of hyperparameters, such as the number of hidden layers, the number of nodes per layer, the kind of activation function, and the learning rate, greatly influence how well DL models perform [15]. The traditional methods to tune these hyperparameters, such as grid search or random search, have significant drawbacks [12]. They can be computationally expensive and inefficient in exploring the hyperparameter space, and they do not consider the interactions between hyperparameters, which may impact the model’s performance [16]. On the other hand, Bayesian Optimization (BO) is a compelling solution, offering a conscientious and efficient approach for hyperparameter tuning in DL models [16]. This probabilistic model-based optimization technique iteratively updates the belief about the objective function to find the maximum value in a minimum number of steps [15, 16].

In this study, we explore the efficacy of Bayesian Optimization in fine-tuning hyperparameters for a deep-structured learning model dedicated to classifying students’ confusion levels from EEG data. Confusion assessment is particularly challenging due to the subtle nature of the cognitive states of the brain signal. The proposed optimal DL model for the ODL-BCI maps hyperparameters to a probability score on the objective function, creating a probabilistic and accurate function model. Unlike traditional methods, it uses information from past evaluations for future selection, thus offering an efficient exploration of the hyperparameter space. By incorporating the complex interaction of hyperparameters and balancing exploration and exploitation in the hyperparameter space, ODL-BCI improves the performance of the DL model, which surpasses traditional ML and state-of-the-art approaches in accuracy and efficiency.

Our research contributes to the field by:

- Presenting a DL model optimized by Bayesian techniques for EEG brain data interpretation.
- Empirically validating the model against conventional classifiers and stateof-the-art methods.
- Establishing the groundwork for real-time BCI applications in educational settings to monitor and enhance student engagement.

The ramifications of this work extend beyond the educational sphere, suggesting potential adaptations in neurotherapeutic settings and human-computer inter-action paradigms.

The rest of this paper is organized as Section 2 literature review, a summary of similar existing works. Then, we move to Section 3 to explain the experimental setup and how we got EEG data. In Section 4, we detail our proposed DL method. We share the results of our experiments in Section 5 and then wrap up with Section 7, where we conclude and discuss future works, suggesting where this research could go next.

## 2. Related Work

The application of DL and ML models explored several studies in braincomputer interfaces (BCIs) for various cognitive tasks. To predict motor imagery tasks from multi-channel EEG data, Miah et al. [1] present the ensemble method called CluSem based on clustering. Their approach achieves high accuracy in classifying motor imagery tasks, showcasing its potential for improving the performance of BCIs. The study contributes to developing more accurate and reliable brain signal analysis techniques. Kabir et al. [16] conduct a comprehensive performance analysis of dimensionality reduction algorithms in ML models for cancer prediction. Their results clarify how various dimensionality reduction methods affect cancer prediction models’ predictive accuracy. The present study significantly contributes to ML model optimization for precise cancer diagnosis and prediction.

Santamaŕıa-Vázquez et al. [11] propose a robust asynchronous control method for ERP-based BCIs using DL. Their research focuses on enhancing the usability and effectiveness of BCIs by leveraging DL techniques. The study demonstrates the potential of DL for achieving reliable and efficient asynchronous control of BCIs, facilitating improved communication between humans and machines. Wu et al. [15] the authors propose an automated approach for hyperparameter tuning in deep learning-based side-channel analysis. The authors demonstrate improved performance in analyzing side-channel vulnerabilities by optimizing the hyperparameters. This research contributes to advancing automated techniques for enhancing the effectiveness of deep learning-based side-channel analysis.

Men and Li [7] investigate using EEG signals to detect student confusion in Massive Open Online Courses (MOOCs). Their study demonstrates the potential of EEG-based analysis in monitoring and understanding the cognitive states of students during online learning. By detecting confusion, this research contributes to developing effective interventions and personalized learning strategies in MOOCs, ultimately enhancing the learning experience. Kunang et al. [13] the authors investigate the application of DL and hyperparameter optimization techniques for classifying attacks in an intrusion detection system (IDS). The study demonstrates the effectiveness of DL models in accurately classifying different types of attacks and the importance of hyperparameter optimization in improving the performance of the IDS. This research contributes to cybersecurity by showcasing the potential of DL and hyperparameter optimization for robust attack classification in IDS.

Cooney et al. [12] evaluate the impact of hyperparameter optimization in the machine and DL methods for decoding imagined speech using EEG signals. Their research emphasizes the importance of optimizing hyperparameters for accurately and reliably decoding imagined speech from EEG data. The study provides valuable insights into optimizing ML models for understanding speech-related brain activity. Reñosa et al. [17] focus on classifying confusion levels using EEG data and artificial neural networks (ANNs). Their study demonstrates the potential of EEG-based classification in assessing cognitive states, such as confusion. By leveraging ANNs, the authors provide insights into using EEG data for real-time cognitive assessment, with potential applications in various fields, including education and human-computer interaction.

Miah et al. [18] using brain-machine interfaces (BMIs) to forecast the orientations of voluntary hand movements and develop a real-time EEG classification system. Their study demonstrates the feasibility of using EEG signals to accurately classify hand movement directions in real time, contributing to the advancement of BMIs. The research showcases the potential of EEG-based BMIs for applications in prosthetics, rehabilitation, and assistive technology. Thomas et al. [19] examine how DL methods are used in BCIs. According to their research, convolutional neural networks (CNNs) and recurrent neural networks (RNNs), two DL algorithms, perform better at accurately classifying brain signals than more conventional techniques. This demonstrates how DL can improve the functionality and performance of BCIs, opening the door to more efficient brain-computer communication.

While these existing papers have significantly contributed to the BCIs field, our paper addresses notable gaps. Firstly, many current approaches need more focus on hyperparameter optimization, relying on traditional methods that might limit BCI model optimization. Our paper introduces more efficient Bayesian optimization for hyperparameter tuning. Secondly, there is a need for more specialized models for classifying cognitive states, such as confusion levels, using EEG signals. Our paper bridges this gap by introducing ODL-BCI, designed explicitly for personalized education and cognitive state monitoring. Thirdly, the efficiency required for practical BCI deployment in real-world scenarios should be considered. Our paper emphasizes both effectiveness and efficiency. Lastly, while previous studies primarily report accuracy, our comprehensive analysis of sensitivity, precision, and F-score demonstrates the practical advantages of ODL-BCI over existing algorithms, highlighting its novelty in these critical aspects.

## 3. Experimental Paradigm and Signal Acquisitions

### 3.1. NeuroSky MindSet

The NeuroSky MindSet EEG headset is used by Wang et al. [20] to record the “confused student EEG brainwave” dataset. This headset is an innovative BCI device engineered for the measurement and recording of electrical brain activity through electroencephalography (EEG) [21]. It is primarily marketed as a BCI device, allowing users to interact with various applications and devices using their brainwave activity [22]. The main feature of the NeuroSky MindSet headset is equipped with an EEG sensor, which is its primary component for measuring the brain’s electrical activity [23]. This single dry electrode is strategically positioned on the FP1 area of the forehead to capture brainwave data. Complementing this, the headset also employs reference and ground sensors located on the ear clip [21]. These are essential for establishing a baseline for the EEG measurements, ensuring the captured data remains accurate and devoid of external interference [24]. It records numeric values determined by proprietary algorithms, reflecting the user’s mental states like attention and meditation. The device also logs numeric values for frequency bands, capturing data from 0 to 60Hz every half-second [17].

It has 14 metallic electrodes mounted on a plastic base, meticulously positioned on the scalp in accordance with the internationally recognized 10-20 system [25]. Refer to Figure 1 for a visual representation of the electrode layout, aligning with the 10-20 system which is used by Wang et al. [20]. The 10-20 system’s meticulous electrode placement ensures consistency and standardization by distributing electrodes based on precise percentages of the scalp’s left-right and front-back dimensions [26]. This adherence to the 10-20 system simplifies integration with existing EEG research and data analysis methodologies [5].

**Figure 1:**
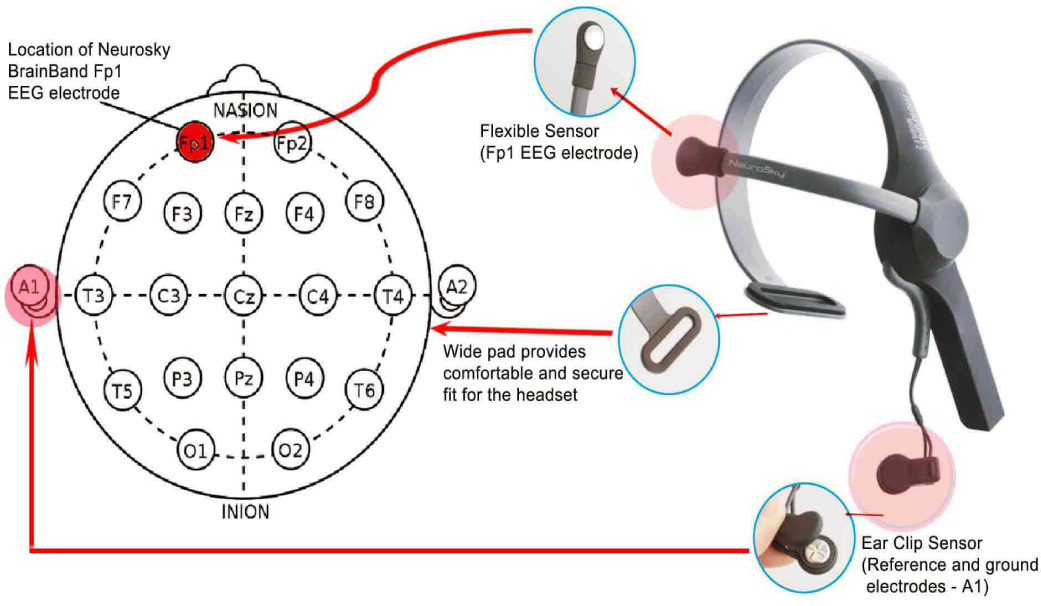
NeuroSky headset electrodes distribution according to the 10-20 international system. Source: NeuroSky.

### 3.2. EEG Dataset Background and Description

The primary objective of the EEG dataset was to investigate the relationship between students’ confusion levels and EEG signals while watching Massive Open Online Courses (MOOC). The dataset was inspired by a pilot study conducted by Wang et al. [20], where college students’ EEG signals were collected to determine their confusion levels when exposed to MOOC content.

Ten college students were enlisted in this dataset to participate in the study. Each student is equipped with a wireless single-channel Neurosky MindSet EEG headset. [27], which precisely measured cerebral activity over the frontal lobe. These students were tasked with watching a set of ten 2-minute-long videos. The MindSet EEG device adeptly extracted a comprehensive set of features, as outlined in Table 1, to capture various aspects of cognitive responses. Notable features included “Attention,” serving as a metric for the students’ mental focus, and “Meditation,” quantifying levels of calmness.

**Table 1:** Features extracted from EEG NeuroSky MindSet.

| Features | Description | Sampling Rate |
| --- | --- | --- |
| Attention | Proprietary measure of mental focus | 1 Hz |
| Meditation | Proprietary measure of calmness | 1 Hz |
| Raw | Raw EEG signal | 512 Hz |
| Delta ($\delta$) | 1-3 Hz of power spectrum | 8 Hz |
| Theta ($\theta$) | 4-7 Hz of power spectrum | 8 Hz |
| Alpha1 ($\alpha$) | Lower 8-11 Hz of power spectrum | 8 Hz |
| Alpha2 ($\alpha$) | Higher 8-11 Hz of power spectrum | 8 Hz |
| Beta1 ($\beta$) | Lower 12-29 Hz of power spectrum | 8 Hz |
| Beta2 ($\beta$) | Higher 12-29 Hz of power spectrum | 8 Hz |
| Gamma1 ($\gamma$) | Lower 30-100 Hz of power spectrum | 8 Hz |
| Gamma2 ($\gamma$) | Higher 30-100 Hz of power spectrum | 8 Hz |

Furthermore, the original EEG signals were averaged and represented by the “Raw” feature. Features from various power spectrum frequency regions were also included in the dataset. These features were sampled at a frequency of 2 Hz, providing a detailed insight into students’ cognitive states. The total number of data samples for this selected dataset is 12,811, where confused are 6,567 and not-confused are 6,244. Based on the duration of the material presented to each student, these data samples are split into average values of 120. With ten data points from each student, this yields 100 data points. The researchers cut out the first and last 30 seconds from each visual material to represent roughly 60 seconds of data samples in each data point. Only the middle portion of each recording was shown to the participating students in the experiment where the dataset was gathered. So, each visual material had an approximate duration of two minutes or less.

Figure 2 shows the extracted features’ correlation matrix with a color bar that emphasizes the likelihood that each extracted feature will be correlated with the other. There is a chance that the diagonal will have one (off-white). As can be seen, the hierarchical clustering approach was used to display the correlation matrix. Light colors indicate positive correlations, while deep shades indicate negative correlations. The box shows the values of the correlation coefficient, either positive or negative. The colors are mapped with varying coefficient values on the right side. As can be seen, the correlation between attention and meditation and the remaining features is inverse, and the highest linear correlation was found between the extracted features’ beta and gamma signals. The ODL-BCI datasets are available for anyone athttps://github.com/MdOchiuddinMiah/ODL-BCI/tree/main/Datasets.

**Figure 2:**
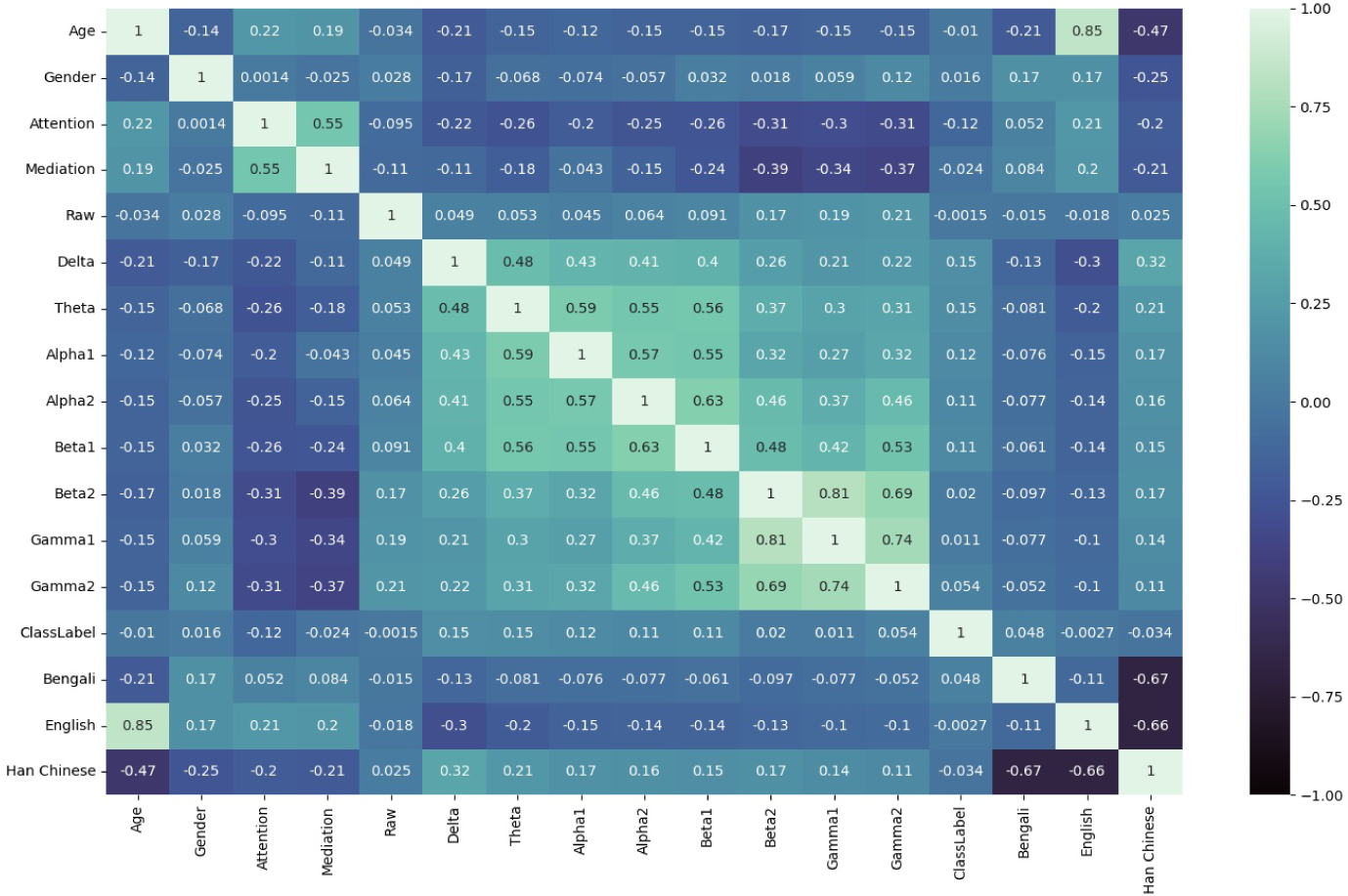
Correlation matrix plot of the extracted features from EEG neuroheadset.

## 4. Classification Techniques

Classification techniques play a pivotal role in the realm of BCIs to analyze EEG data [1]. The application of these techniques aids in classifying different mental states based on the acquired data [18, 5]. In this study, we utilized several baseline models, including Decision Tree, AdaBoost, Bagging, Multilayer Perceptron (MLP), Näıve Bayes, Random Forest, Support Vector Machines (SVM), XG Boost, and a specially designed DL model. Each of these classifiers offers different strengths, and their performance tends to vary depending on the nature of the dataset and problem at hand [28]. Our objective was to explore and compare the efficiency of these classifiers on the EEG data, with particular emphasis on our proposed DL model.

### 4.1. Baseline Models

This section presents an overview of the baseline models employed in our investigation. These models serve as essential benchmarks for assessing the performance of our proposed DL model, *ODL − BCI*, in classifying students’ confusion levels from EEG data.

#### 4.1.1. Decision Tree

Decision trees are renowned for their interpretability and simplicity [18]. They partition the dataset based on the most relevant features, constructing a tree-like structure that aids decision-making [1]. Here, we implement the *C*4.5 [29] algorithm.

#### 4.1.2. AdaBoost

AdaBoost is an ensemble learning method that combines several weak classifiers’ predictions to produce a robust classifier[1]. It focuses on rectifying the errors of prior classifiers, ultimately enhancing classification accuracy [28]. We used the *C*4.5 algorithm and 100 learners to train and build a robust classifier.

#### 4.1.3. Bagging

Bagging, or Bootstrap Aggregating, is another ensemble method that crafts numerous base models using random subsets of the dataset [5]. The amalgamation of these models results in better predictions, reducing overfitting [1]. We used *DecisionTreeClassifier* as the base estimator.

#### 4.1.4. Multilayer Perceptron (MLP)

MLP, an artificial neural network, consists of multiple interconnected layers [1]. It excels at capturing complex patterns in data [1]. Here, we deploy MLP to gauge its performance relative to our DL approach. We used 100 hidden layers, activation function *relu*, *adam* for weight optimization, and learning rate 0.0001.

#### 4.1.5. Näıve Bayes

The Näıve Bayes classifier relies on Bayes’ theorem and feature independence assumption [5]. Despite its simplicity, it demonstrates surprising effectiveness, particularly in text classification tasks [28]. We used the *MultinomialNB* classifier to construct the model.

#### 4.1.6. Random Forest

Random Forest is an ensemble approach that builds multiple decision trees during training and aggregates their results[28]. Its robustness and suitability for high-dimensional data make it a valuable baseline [28]. We used 100 trees in the forest with *gini* classifier and minimum samples 2 to split an internal node.

#### 4.1.7. Support Vector Machines (SVM)

SVMs are potent classifiers that aim to identify a hyperplane best suited to segregate data points from different classes [18]. They shine in high-dimensional spaces and complex datasets [28]. We implement SVM with *linearkernel* and *predict proba* to enable probability estimation.

#### 4.1.8. XG Boost

Renowned for its speed and performance, XG Boost is a gradient-boosting algorithm that builds an ensemble of decision trees iteratively, focusing on reducing predictive errors [30]. We used *binary* : *logistic* for binary classification with logistic regression [30] and 100 decision trees in the ensemble with one random seed for reproducibility.

### 4.2. Proposed Deep Learning Model

Our proposed method incorporates an optimal deep-learning model with hyperparameters optimized by Bayesian optimization. It has been tailored explicitly for BCI data analysis. The below sections provide a detailed outline of the model structure, parameter selection, optimization technique, model construction, and training.

#### 4.2.1. Model Structure and Parameter Selection

The architecture of our DL model consists of an output layer, multiple hidden layers, and an input layer. The quantity of input features and output classes determines the nodes in the input and output layers. The architecture and parameters of the hidden layers are meticulously selected through the Bayesian optimization process, as highlighted in Table 2 and 3. The model features four hidden layers (H1, H2, H3, H4) with varying numbers of nodes (200, 100, 50, 16, respectively). The activation functions for these layers, selected based on their performance during the optimization process, are either Rectified Linear Units (RELU) or Parametric Rectified Linear Units (PRELU). The learning rates for each layer are set at 0.001 or 0.01, again based on the optimization process. Binary crossentropy as loss function and optimizer Adam used to train the model. Two nodes on the output layer represent the class label Confused and Not-Confused, which used the activation function Sigmoid to calculate the output of the nodes.

**Table 2:** Optimized ODL-BCI hyperparameters.

| Hyperparameters | List of Values (N) |
| --- | --- |
| No. of Layers | $1 \leq N \leq 6$ |
| Nodes per Layer | $4 \leq N \leq 200$ |
| Activation Function | Sigmoid, RELU, PRELU, ELU, Swish, Tanh |
| Learning Rate | 0.000001, 0.00001, 0.0001, 0.001, 0.01, 0.1 |

**Table 3:** Summary of the Best Parameters.

|  | Grid Search | Bayesian Algorithm |
| --- | --- | --- |
| Number of hidden Layer | 3 | 4 |
| Size of hidden Layers | 150-60-20 | 200-100-50-16 |
| Activation Function | PRELU, Sigmoid | RELU, PRELU, Sigmoid |
| Learning Rate | 0.001 | 0.001 |
| Time [h:m:s] | 23:10:40 | 12:10:20 |
| Number Of runs | 42,552 | 21,200 |
| Accuracy | 68% | 74% |

#### 4.2.2. Bayesian Optimization

The centerpiece of our hyperparameter optimization process is the implementation of Bayesian Optimization, as described in Algorithm 2. Initially, the hyperparameters are set to none. Each iteration includes sampling parameters from the search space, performing cross-validation, calculating an evaluation metric, and adding the parameters and evaluation metric to the hyperparameters set. This process continues until the top hyperparameters that yield the highest evaluation metric are obtained.

#### 4.2.3. Model Construction and Training

Our DL model’s creation and training process are governed by the procedures listed in Algorithm 1. The model starts by dividing the BCI data into 80% training and the remaining 20% test datasets. Following this, the Bayesian optimization process generates a set of hyperparameters. The model is initialized using these parameters, trained using the training data, and evaluated on the test data. This cycle is repeated for each set of hyperparameters, with the most accurate model on the test data identified as the best model. Its hyperparameters are then stored.

The developed DL model, with its bespoke structure and optimization pro-cess, offers a potent approach to BCI data analysis. As confirmed by the results shown in Table 5, the model surpasses traditional ML classifiers, demonstrating an accuracy of 74%. This underscores the viability and superiority of DL models optimized with Bayesian optimization in BCI applications. The ODL-BCI implementation with compared models source codes can be accessed by anyone athttps://github.com/MdOchiuddinMiah/ODL-BCI/tree/main/Algorithms.

## 5. Experiments

We performed a series of experiments to test the effectiveness of our proposed DL model. Our experimental setup, the optimized hyperparameters of the model, and the results obtained will be discussed in the following sections.

### 5.1. Experimental Setup

All experiments were performed on a MacBook Pro with an Intel Core i7 3.3 GHz Dual-Core processor and 16 GB of RAM. Python 3.8 was utilized as the primary language for coding, while the Scikit-learn 0.21.2 and TensorFlow 2.9 libraries facilitated the implementation of the machine-learning models. Spyder 3.3.1 (https://www.spyder-ide.org), a Python development environment, was used for coding and running the models.

Using popular ML algorithms as benchmarks, our model’s performance is evaluated using classification metrics, including accuracy, precision, recall, and F-score. Eq. 1 is used to quantify the accuracy; when *x_i_* is correctly classified, *assess*(*x_i_*) = 1; when *x_i_* is misclassified, *assess*(*x_i_*) = 0 [1]. The computations are displayed in Eq. 2 to 4 [28, 18]. The weighted average values of precision, recall, and F-score are considered. The true positive rate (TPR) and false positive rate (FPR) trade-off between the classifiers was visualized and quantified using ROC curves and AUC scores. The calculations are defined in Eqs. 5 and 6 [18]. The true positive rate (TPR) is plotted against the false positive rate (FPR) in the ROC curve. The lowest threshold is represented by a line, *y* = *x*, in the au-ROC curve, where correctly classified data points indicate 1 and misclassified instances are revealed as 0 [28].

#### Algorithm 1 Proposed deep learning model with optimized hyperparameters by Bayesian Optimization

**Input:** BCI data *D*

**Output:** Optimized deep learning model

**Method:**

1. Split *D* into *train data* and *test data*;
2. hyperparams = BayesianOptimization();
3. best accuracy = 0;
4. best model = None;
5. best hyperparams = None;
6. **for** params in hyperparams **do**
7. nl = params[’no of layers’];
8. npl = params[’nodes per layer’];
9. af = params[’activation function’];
10. lr = params[’learning rate’];
11. model = initialize model(nl, npl, af, lr);
12. train model with *train data*;
13. accuracy = evaluate model(*test data*);
14. **if** accuracy > best accuracy **then**
15. best accuracy = accuracy;
16. best model = model;
17. best hyperparams = params;
18. **end if**
19. **end for**
20. **return** best model with best hyperparams;

#### Algorithm 2 Bayesian Optimization for proposed deep learning model ODL-BCI

**Output:** Optimized hyperparams

**Method**:

1. *hyperparams* = None;
2. **for** each iteration **do**
3. Sample *params* from the search space;
4. Perform cross-validation with *params*;
5. Calculate *evaluation metric*;
6. *hyperparams ∪ {params, evaluation metric}*;
7. end for
8. Sort *hyperparams* in descending by *evaluation metric*;
9. **return** top *hyperparams*;

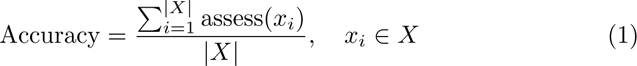

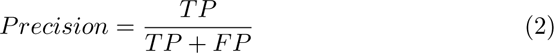

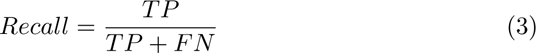

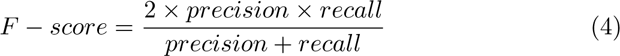

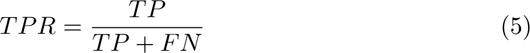

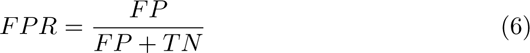

The notations TP, TN, FP, and FN denote the true positive, true negative, false positive, and false negative results, respectively [1].

### 5.2. Results

Our empirical results are embodied in Tables 2 through 5, which jointly provide a comprehensive view of the optimal DL model’s hyperparameter tuning, comparative performance, and preferred configuration.

We initially evaluated the performances of the ODL-BCI model with different learning parameters. These parameters were tested using a comprehensive hyperparameter list, including the number of layers, nodes per layer, activation function types, and learning rate, as Table. 2 detailed.

Table 3 tabulates the best parameters suggested by the grid search and Bayesian optimization algorithms. The total execution time taken by the Grid Search is more than 23 hours, almost double that of the Bayesian Algorithm. Also, the number of runs for the Bayesian Algorithm is half the number for Grid Search. A primary concern is the accuracy, and our observations indicate that Bayesian optimization outperforms grid search regarding model accuracy 74% and time efficiency.

The optimal hyperparameters for the DL model on students’ confusion EEG dataset were selected based on the best performance metrics and are illustrated in Table 4. This table elucidates the most practical combination of hidden layers, nodes per layer, activation functions, and learning rate for the proposed model. A four-layer model is proposed, employing RELU and PRELU activation functions and adopting learning rates of 0.001 and 0.01.

**Table 4:** The best performed hyperparameters for the deep structured model on students confusion EEG dataset.

| Hidden Layers | Nodes per Layer | Activation Function | Learning Rate |
| --- | --- | --- | --- |
| Layer $H_1$ | 200 | RELU | 0.001 |
| Layer $H_2$ | 100 | RELU | 0.001 |
| Layer $H_3$ | 50 | PRELU | 0.01 |
| Layer $H_4$ | 16 | PRELU | 0.001 |

**Figure 3:**
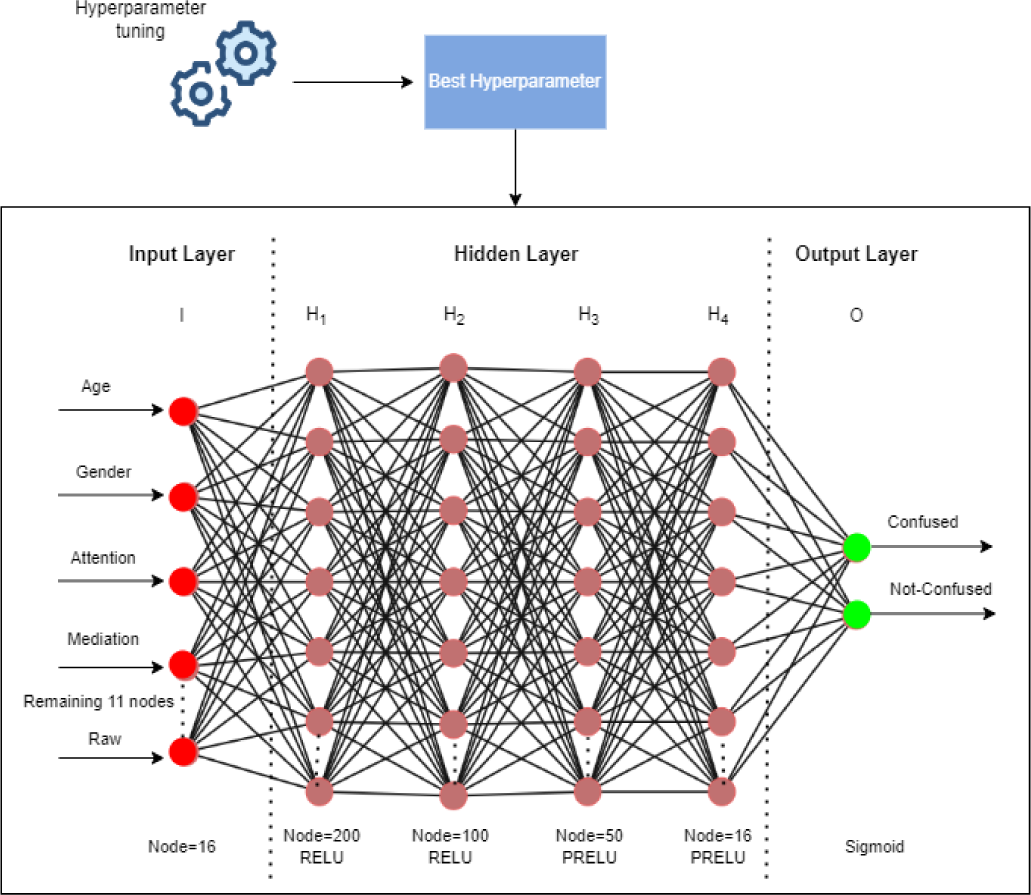
The work-flow of the proposed optimal deep learning model for student’s confusion EEG dataset.

Next, the performance of the ODL-BCI model was compared with existing methods, such as Decision Tree, AdaBoost, Bagging, MLP, Näıve Bayes, Random Forest, SVM, and XG Boost. The comparative analysis of accuracy, precision, recall, and F-score of these classifiers on students’ confusion level EEG data is presented in Table 5. The proposed DL model significantly outperforms the existing classifiers. The ODL-BCI model achieves 74% accuracy, precision, recall, and F-score, demonstrating superior performance to the other tested classifiers. While existing ensemble approaches can reach up to 69.0% accuracy, single classifiers in this dataset failed to achieve more than 60% accuracy. The table indicates that ODL-BCI outperforms other ensemble classifiers used for this task by more than 5.0%. Fig. 4 displays the decision boundaries of the suggested method compared to the classifiers used for this dataset.

**Table 5:** Accuracy and Average Sensitivity/Recall, Precision and F-score on EEG confusion dataset.

| Classifiers | Accuracy (%) | Precision | Recall | F-score |
| --- | --- | --- | --- | --- |
| Decision Tree | 60 | 0.60 | 0.60 | 0.60 |
| AdaBoost | 61 | 0.61 | 0.61 | 0.61 |
| Bagging | 64 | 0.65 | 0.64 | 0.64 |
| MLP | 66 | 0.66 | 0.66 | 0.66 |
| Naïve Bayes | 58 | 0.58 | 0.58 | 0.58 |
| Random Forest | 69 | 0.69 | 0.69 | 0.69 |
| SVM | 58 | 0.58 | 0.58 | 0.58 |
| XG Boost | 67 | 0.67 | 0.67 | 0.67 |
| **ODL-BCI** | **74** | **0.74** | **0.74** | **0.74** |

**Figure 4:**
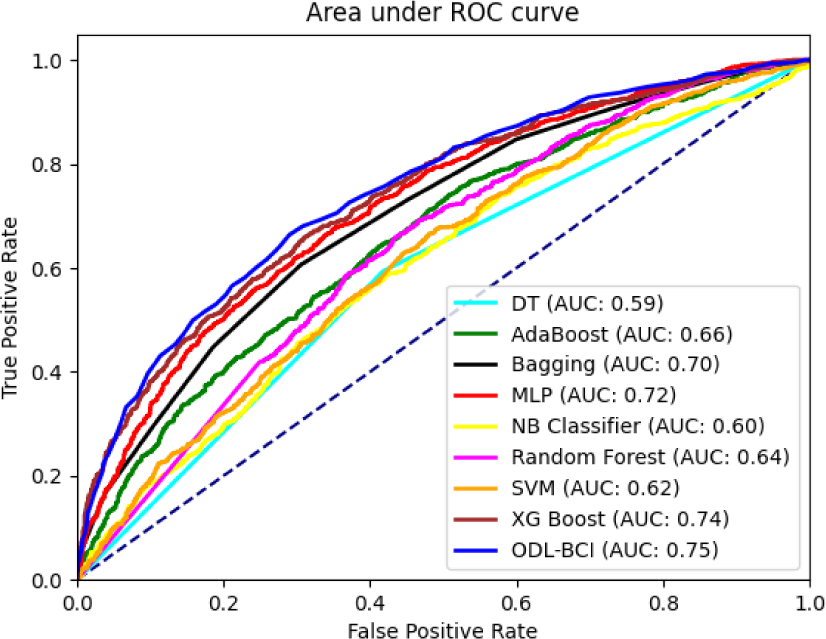
ROC and AUC analysis of ODL-BCI model compared to popular classifiers on students confusion EEG dataset.

For a more in-depth understanding of our DL model’s behaviour, we monitored its training progress over 2000 epochs. Figures 5 and 6 depict the accuracy and loss over time. Notably, we observed that after approximately 400 epochs, accuracy and loss reached stable levels. The accuracy plateaued at around 0.67, while the loss remained below 0.60. These observations suggest that our model effectively captured the underlying patterns in the data, achieving remarkable performance.

**Figure 5:**
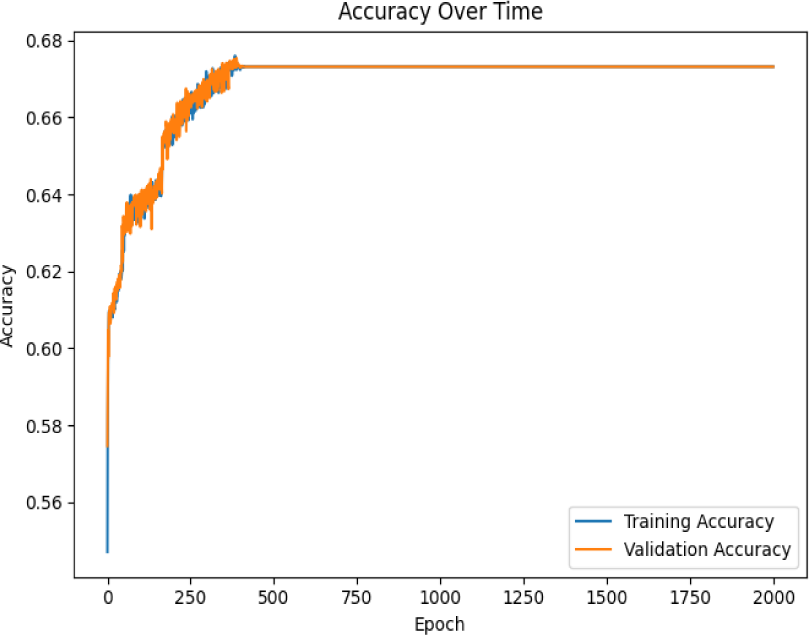
Accuracy over time after 2000 epochs for proposed ODL-BCI model using student’s brain data.

**Figure 6:**
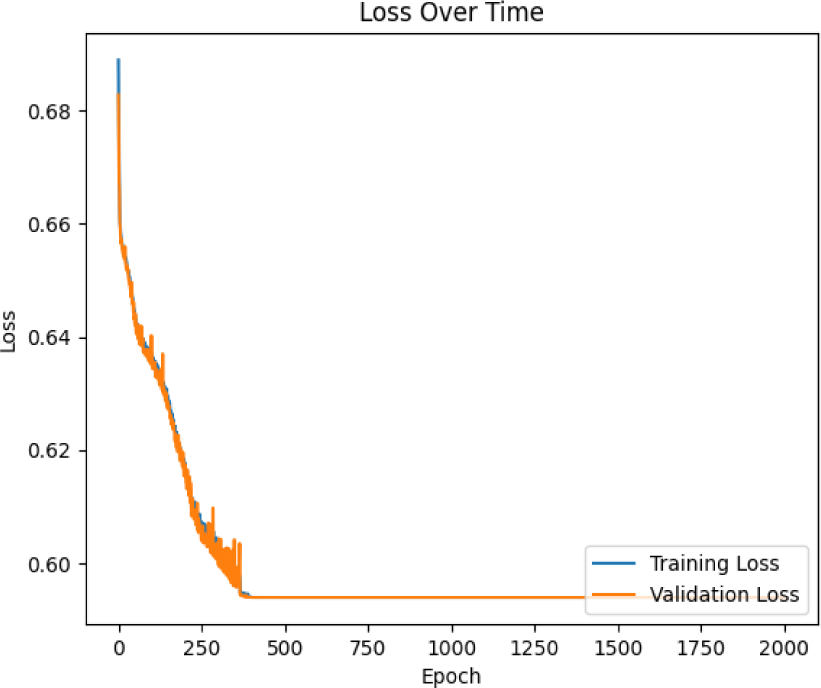
Loss over time for optimized deep learning model after 2000 epochs.

### 5.3. Comparison with State-of-the-Art Methods

To evaluate the prowess of our proposed ODL-BCI model, we compared its performance with several state-of-the-art methods previously employed on the students’ confusion EEG dataset. This comparison sheds light on the effectiveness of our novel approach in pushing the boundaries of classification accuracy.

As illustrated in Table 6, our ODL-BCI model achieved a remarkable accuracy of 74.0%. This accomplishment surpasses the performance of other notable methods. For instance, SVM with a linear kernel, as documented by Ni et al. [31], attained an accuracy of 67.2%. Meanwhile, Bidirectional LSTM and RNN-LSTM, also explored by Ni et al. [31], achieved accuracies of 73.3% and 69.0%, respectively. Haohan et al. [20] presented two approaches—Pre-defined Confusion Level and User-defined Confusion Level—with accuracies of 67.0% and 56.0%, respectively.

**Table 6:** Compare with state-of-the-art methods used students confusion EEG dataset.

| Classification Methods | Accuracy (%) |
| --- | --- |
| SVM (linear kernel) (Ni et al. [31]) | 67.2 |
| Bidirectional LSTM (Ni et al. [31]) | 73.3 |
| RNN-LSTM (Ni et al. [31]) | 69.0 |
| LLM using BERT (Kostas et al. [32]) | 59.0 |
| Pre-defined Confusion Level (Haohan et al. [20]) | 67.0 |
| User-defined Confusion Level (Haohan et al. [20]) | 56.0 |
| **Proposed ODL-BCI** | **74.0** |

In addition to our proposed ODL-BCI model, we explored the application of Large Language Models (LLM) for the classification task, inspired by the work of Kostas et al. [32]. Leveraging the BERT (Bidirectional Encoder Representations from Transformers) model for sequence classification, our implementation achieved an accuracy of 59%. The LLM approach involves tokenizing the EEG data into sequences and training a transformer-based model on the provided dataset. While our primary focus remains on the ODL-BCI model, utilizing LLM presents an alternative perspective on EEG-based classification tasks. The comparative performance of the ODL-BCI model and the LLM approach is detailed in Table 6, highlighting the nuanced outcomes and shedding light on the potential impact of different methodologies in confusion level detection.

Our ODL-BCI model’s substantial superiority over these state-of-the-art methods is evident, marking a significant advancement in EEG-based confusion level classification.

## 6. Threats to Validity

In discussing the findings of our study, it is essential to acknowledge potential threats to validity that may impact the robustness and generalizability of our results. One potential threat is related to the dataset used in our experiments. While we carefully curated a dataset of EEG data to represent students’ confusion levels, the inherent variability in individual cognitive responses may introduce bias. Additionally, the relatively modest size of our dataset poses challenges in achieving complete generalizability. Furthermore, as with any machine learning model, the performance of our proposed ODL-BCI is contingent on the chosen hyperparameters and model architecture. Although we employed rigorous optimization techniques such as Bayesian Optimization, the sensitivity of these choices to different datasets and tasks remains a consideration. Lastly, using a specific preprocessing pipeline and feature extraction methods may impact the reproducibility of our results in different experimental setups. Addressing these challenges and exploring their potential impact on the outcomes of our study is crucial for a comprehensive understanding of the proposed model’s applicability.

## 7. Conclusions & Future Work

This study introduced Bayesian optimization as a robust methodology for fine-tuning hyperparameters in deep learning-based BCI models, aiming to classify students’ confusion levels using EEG data. We developed a tailored framework that combines the power of DL with Bayesian optimization techniques, creating a model referred to as ODL-BCI.

Our findings demonstrate that Bayesian optimization is exceptionally effective for optimizing hyperparameters in EEG-based cognitive state classification. The ODL-BCI model, enriched with Bayesian optimization, outperformed conventional ML classifiers and even state-of-the-art methods on the “Confused student EEG brainwave data” dataset. This model achieved an impressive accuracy of 74 percent, underscoring its potential as a valuable tool in the educational sector for real-time confusion level assessment.

Although the proposed model shows promising results, future work can further enhance its capabilities. For instance, the model can be tested with a larger dataset to verify its scalability. Incorporating other types of neural networks and more advanced optimization algorithms can also be explored to improve the model’s performance. Furthermore, real-world deployment of the model in the educational sector can be examined to assess its practical utility in enhancing student confusion and learning outcomes.

